# “Anthropogenic Threats and Conservation Practices: Evaluating LULC Changes in Relocated Villages of Panna Tiger Reserve, Madhya Pradesh, India”

**DOI:** 10.1101/2023.03.08.531715

**Authors:** Susmita Khan

## Abstract

Protected areas globally, including India, face anthropogenic threats despite environmental and forest protection laws. To reduce these pressures, conservation related settlement evacuation from protected areas has been a common conservation practice for decades in many countries. Controversies aside, studies have shown that evacuation of settlements from protected areas has a positive impact on forest cover change and biodiversity increase. Here in this study, we conducted this study for the 14 relocated village sites of Panna Tiger Reserve to see the changes in major Land use Land cover changes over 15 years (2004-2019). We observed a significant increase in forested areas in those regions over time, as well as prominent changes in different land cover classes. The increase in Grassland, Open Forest, and Dense Forest from 2004 to 2019 was significant, and settlements in relocated areas were mainly converted to different forest types. Additionally, fieldwork and camera trap data showed that the relocated sites with established woodlands and other forest classes supported higher animal diversity and species composition compared to other sites. This study provides a knowledge base for conservation and management practices by identifying significant changes in various land use and land cover classes.

## 1. Introduction

Nearly 12% of the world’s land is protected by Nature reserves, of which merely 2% comes under biodiversity-rich tropical forests (Rodrigues et al., 2004). Protected Areas (PAs) are essential for maintaining habitat integrity and species diversity (Geldmann et al., 2013). A high density of the human population near and within the PAs causes high anthropogenic pressure on forests and animals (Karanth, 2002; Muller & Zeller, 2002; Sanderson & Redford, 2003; Peh et al., 2005). In India, almost 5 million people live inside PAs with densities exceeding 300 people/Km^2^ and are directly dependent on forest resources (Kutty & Kothari, 2001; Rodgers et al., 2003). Although there are laws (i.e., Indian Wild Life Protection act, 1972) that prohibit hunting, fishing, livestock grazing, and other activities within PAs in India, these practices are still common. They can lead to conflicts with wildlife, crop damage, and livestock and human life loss. (Kothari et al., 1989; Madhusudan & Karanth, 2000; Karanth, 2002; Madhusudan & Mishra, 2003; Treves & Karanth, 2003; Madhusudan, 2004; Barve et al., 2005; Das et al., 2006; Karanth et al., 2006; Kumar & Shahabuddin, 2005).

Conservation-related emigration is widely practiced in Africa, South, and Southeast Asia, and in North America (Brockington & Igoe, 2006), and there are also criticisms of these efforts due to their perceived negative impacts on people who have been relocated (Sato, 2000; Brechin et al., 2002; Wilshusen *et al*., 2002; Brockington, 2003; Chatty & Colchester, 2003; Schmidt-Soltau, 2003; Brosius, 2004; Rangarajan & Shahabuddin, 2006). The Indian government practice settlement relocations in many PAs, and various restrictions have been made for the resource utilization by local people within these PAs (Sharma, 2003; Shahabuddin & Shah, 2003; Kabra, 2006; Rangarajan & Shahabuddin, 2006). Despite controversies, some studies prove the positive impacts of such relocation activities on concerned PAs (Karanth, 2007). It has been found that such areas with evacuation or relocation significantly impact vegetation community restoration over time (Heras, Martin & Espigares, 2008).

In India, most anthropogenic disturbances in and around PA include livestock grazing, fishing, fuelwood and Non-timber Forest Product (NTFP) extraction, and agriculture, causing biodiversity loss, habitat fragmentation, and Landcover changes. (Gadgil & Guha, 1992; Kothari et al., 1995; Murali et al., 1996; Somanathan & Borges, 2000; Rahmani, 2003). These types of anthropogenic pressure ultimately affect ecosystem functioning (Robinson, 1993). Grazing and NTFP extraction may cause a decrease in food and other resources for wild animals leading to crop-raiding and human-wildlife conflict, change in nutrient dynamic, soil erosion and vegetation community alteration causing local extinction of small taxa and increased disease transmission to wild herbivore population (Sekhar, 1998; Madhusudan & Mishra, 2003; Middleton, 2003; Rahmani, 2003). The relocation of villages inside PAs began in the 1960s as a way to promote conservation in India (Rangarajan & Shahabuddin, 2006; Shahabuddin & Shah, 2003; Sharma, 2003; Ganguly, 2004; Shahabuddin et al., 2005; Kabra, 2006; Rangarajan & Shahabuddin, 2006; Karanth & Karanth, 2007). Various restrictions have been placed on the use of PA resources by local people (Sharma, 2003). The number of relocations in India for conservation purposes is less than 1% of the total number of relocations that occur for other reasons (60 million). These relocations can range from forcible to voluntary (Karanth, 2007).

Remote Sensing aids in analyzing the landscape dynamics with the help of satellite images of the earth’s surface acquired at different temporal and spatial scales (NAP, 2008). One of the leading applications of these remotely sensed data is land use Land cover (LULC) change detection by using repetitive coverage of an area (Anderson, 1977; Ingram et al., 1981; Nelson, 1983; Singh, 1984). LULC change detection analysis helps to identify changes in the landscape caused by natural and human factors (CCSP, 2003; Zheng et al., 2009) across varying temporal and spatial scales, representing the confluence of the environmental and human dynamics which govern the transition (Lambin et al., 2003). Panna tiger reserve (PTR) of Madhya Pradesh, India, is one of the 53 tiger reserves of India and was established in 1994. PTR is also one of the biosphere reserves of India. The tiger population of the PTR was eliminated by 2009 because of poaching. Due to the active efforts of tiger translocation, currently, 31±3 tigers are present within the PTR (Jhala et al., 2020).

Village settlements from the core areas of the PTR started in 1987, and out of 17 villages, 14 villages were translocated outside the core areas till 2020 (WII, 2022). Continuous habitat restoration practices in all these village sites showed real success, with the tiger and other species occupying these translocated villages (WII, 2022). However, till now, no studies have been taken to understand the change in LULC patterns in those villages, which can help the PA manager formulate landscape management strategies. Thus, we conducted this study for 14 relocated village sites within PTR’s core region, Madhya Pradesh, India. The objective of the study is to identify the changes in the significant LULC classes over the past 15 years in 14 evacuated village sites of PTR to identify focus areas for grassland management and conservation priorities.

## 2. Materials and methods

### 2.1. Study area

Panna Tiger Reserve (Figure 1) is located in Vindhya Mountain ranges that extend from Southwest to Northeast in the civil districts of Panna, Chattarpur and Damoh of Madhya Pradesh, India, covering an area of 576.13 Km^2^ (Core area) +1021.97 Km^2^ (Buffer area) = 1598.1 Km2. The primary forest type of the PTR is dry deciduous. The area experiences an annual average rainfall of 1118mm, and the average temperature is 24.8°C. A total of 17 villages were situated within the core areas of PTR, of which, between the years 1987 to 2015, a total of 14 villages (Table 1) were relocated to the buffer areas of the PTR. Those 14 village sites are entirely abandoned and now in ruins. Dhodan, Palkoha, and Khariyani-Mynari are the remaining three villages yet to be relocated. PTR and the surrounding territorial forest division of north Panna and south Panna are the only large chunk of wildlife habitat remaining in the fragmented forested landscape of north Madhya Pradesh. It represents one of the critical tiger habitats of central Indian highlands along with its associated species. All analyses were performed within these 14 relocated villages of PTR.

**Figure 1.**
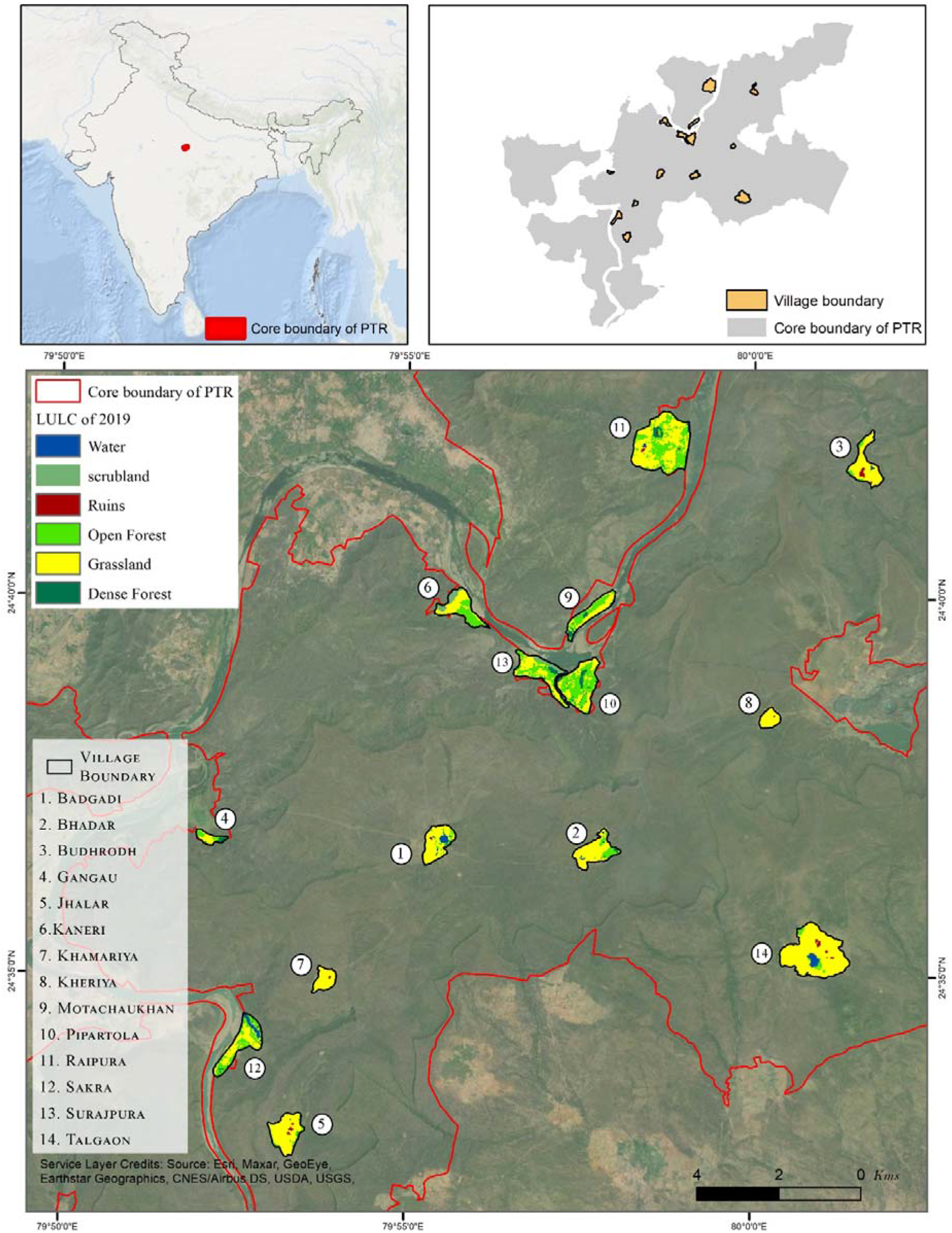
Map of the study area.

**Table 1.** Details of relocated villages of PTR.

| S. No. | Village name | Area (Sq. Km) | Year of relocation<sup>329</sup><br>330 |
| --- | --- | --- | --- |
| 1. | Badgadi | 0.511 | 1987 -91 |
| 2. | Bhadar | 0.536 | 2005-2015 |
| 3. | Budhrodh | 0.561731 | 2008-09 |
| 4. | Gangau | 0.1529 | 2005-2015 |
| 5. | Jhalar | 0.727369 | 2009-10 |
| 6. | Kaneri | 0.589833 | 2007-09 |
| 7. | Khamariya | 0.2467 | 1987 -91 |
| 8. | Kheriya | 0.1869 | 1987 -91 |
| 9. | Motachaukhan | 0.523132 | 2007-09 |
| 10. | Pipartola | 0.96218 | 2007-09 |
| 11. | Raipura | 1.949925 | 2007-09 |
| 12. | Sakra | 0.781733 | 2005-2015 |
| 13. | Surajpura | 0.752077 | 2007-09 |
| 14. | Talgaon | 1.832514 | 2010-13 |

### 2.2. Change detection analysis

#### Data

We downloaded satellite data from the USGS Earth Explorer site. For this study, Landsat images of 30 m resolution were obtained. We used the Landsat 8 satellite image for 2019; and the Landsat 5 satellite image for 2004 due to the unavailability of the Landsat 8 image in 2004 and the inadequate quality of Landsat 7 image. For both years, we used post-monsoon images (i.e., November) for cloud-free images.

#### Image Processing

Satellite processing of the images was performed in ArcGIS 10.7. Atmospheric correction of the images was done using a raster calculator in ArcGIS 10.7. We took six bands of each image for both years. Both images were masked with the village boundaries. These masked files were used for further classification and change detection analysis.

#### Classification

We performed image classification in ERDAS Imagine v14. The two key methods of image classification are Supervised classification and Unsupervised classification. Supervised classification is usually performed when a former field visit has been made, and the classes are assigned depending on the signatures obtained during the field visit. In contrast, unsupervised classification depends on the pixels’ natural spectral reflectance to merge them into the required number of classes.

For this study, we selected supervised classification, as (i) the study area is very small, and (ii) field visits to the sites were made before the analysis. We used eight classes for 2004 classification, i.e., Water(Z1), Ruins (Z2), Dense Forest(Z3), Settlements(Z4), Cropland(Z5), Scrubland(Z6), Open Forest(Z7), Grassland(Z8). By 2019, all the villages were relocated, and settlements and croplands had been changed to other classes. Thus for 2019, we used six classes, i.e., Water (C1), Ruins(C2), Grassland (C3), Open Forest (C4), Dense Forest(C5), Scrubland(C6). Post-supervised classification, we recoded the classified images with thematic codes referencing Google Earth imagery and ERDAS.

#### Change Matrix Generation

Post-recoding, we performed a change detection analysis and generated a Change Matrix. Change matrices help understand the LULC conversion between the various LULC classes considered for the analysis. The change detection analysis was performed using the raster datasets of the LULC maps of 2004 and 2019 in the ERDAS Imagine software. We used the “Matrix Union” tool to generate the matrix between the LULC maps of the years 2004 and 2019.

## 3. Results

### LULC maps

The total classes taken for the LULC maps are Water, Ruins, Dense Forest, Settlements, Cropland, Scrubland, Open Forest, and Grassland. A total of eight classes for 2004 and six classes for 2019 was taken to classify the 14 relocated village sites and to prepare the LULC maps (Figure 2, 3 and 4). Table 2 details the area (in sq. km) occupied by each class and the percentage of land occupied by each LULC type.

**Figure 2.**
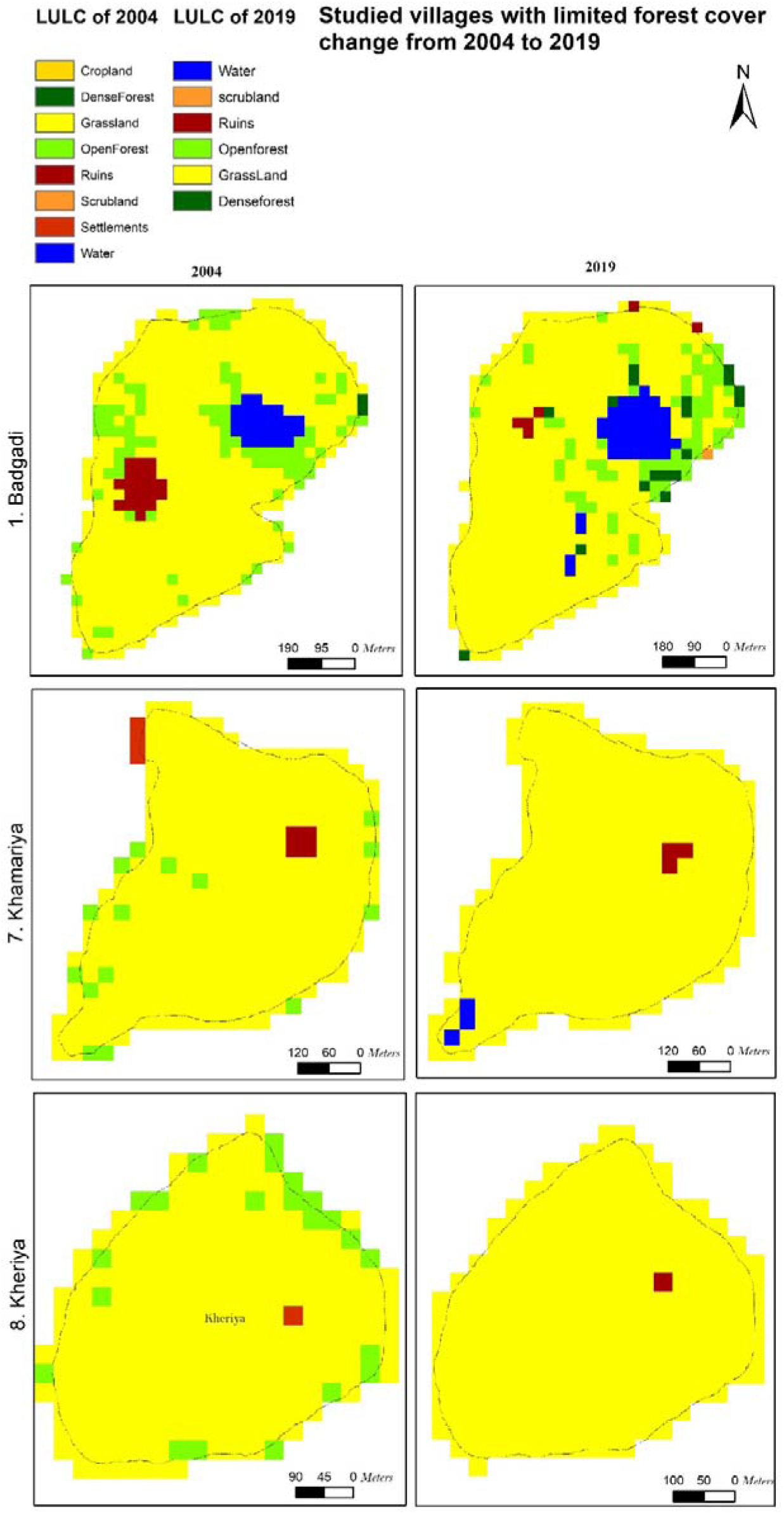
Map of the translocated villages with limited forest cover change from 2004 to 2019

**Figure 3.**
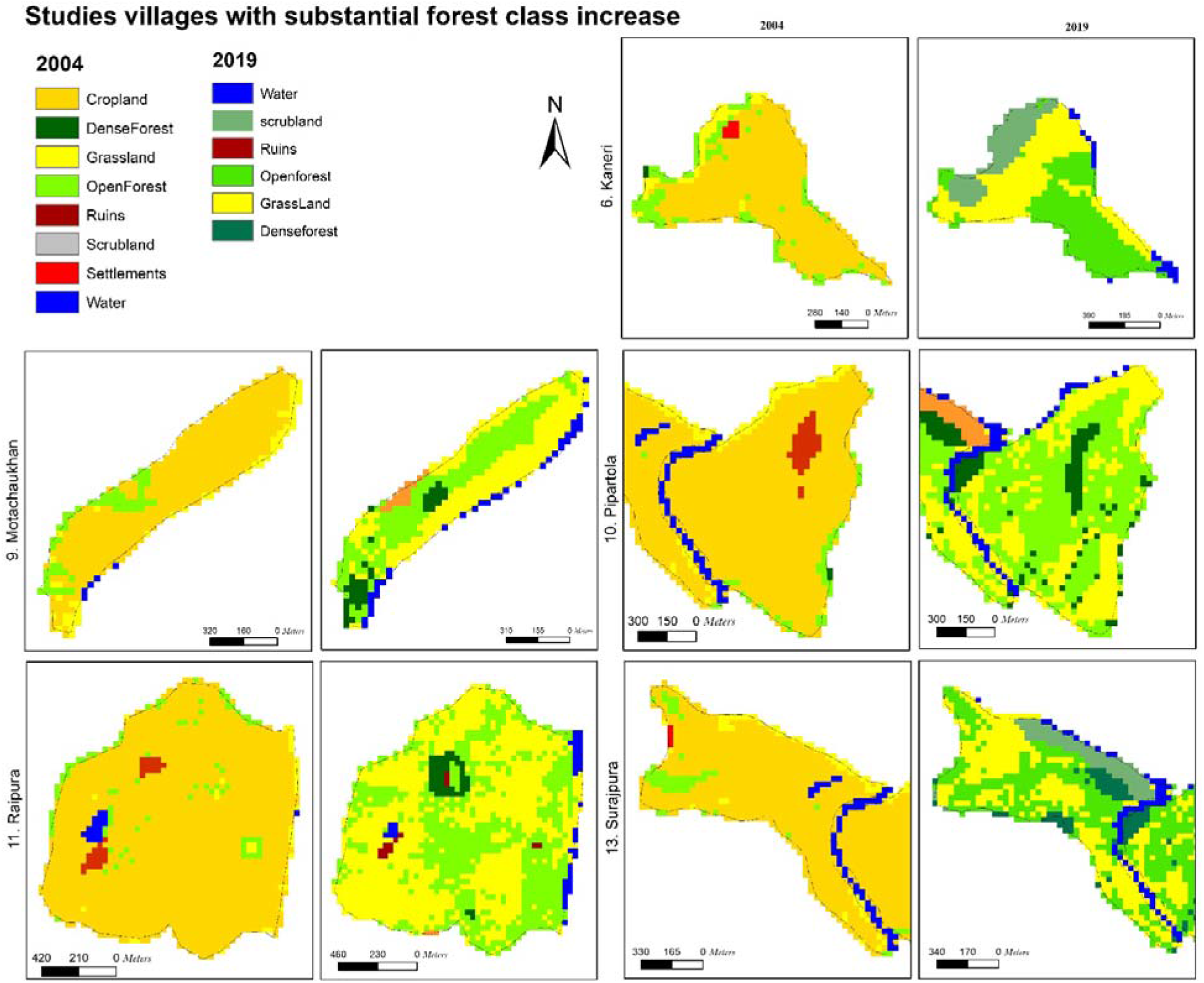
Map of the translocated villages with substantial forest classes change increase from 2004 to 2019.

**Figure 4.**
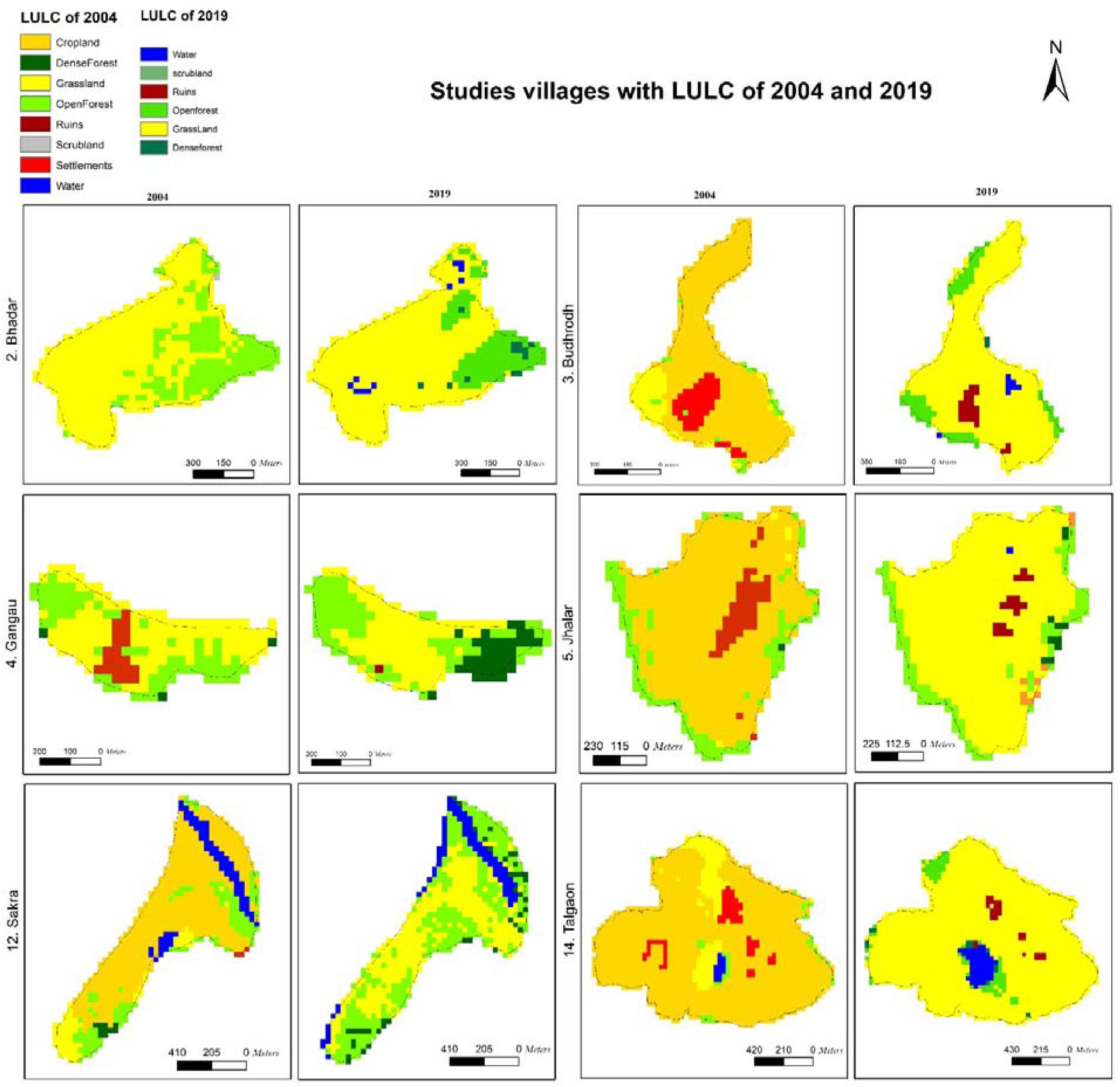
Map of the translocated villages from 2004 to 201

**Table 2.**
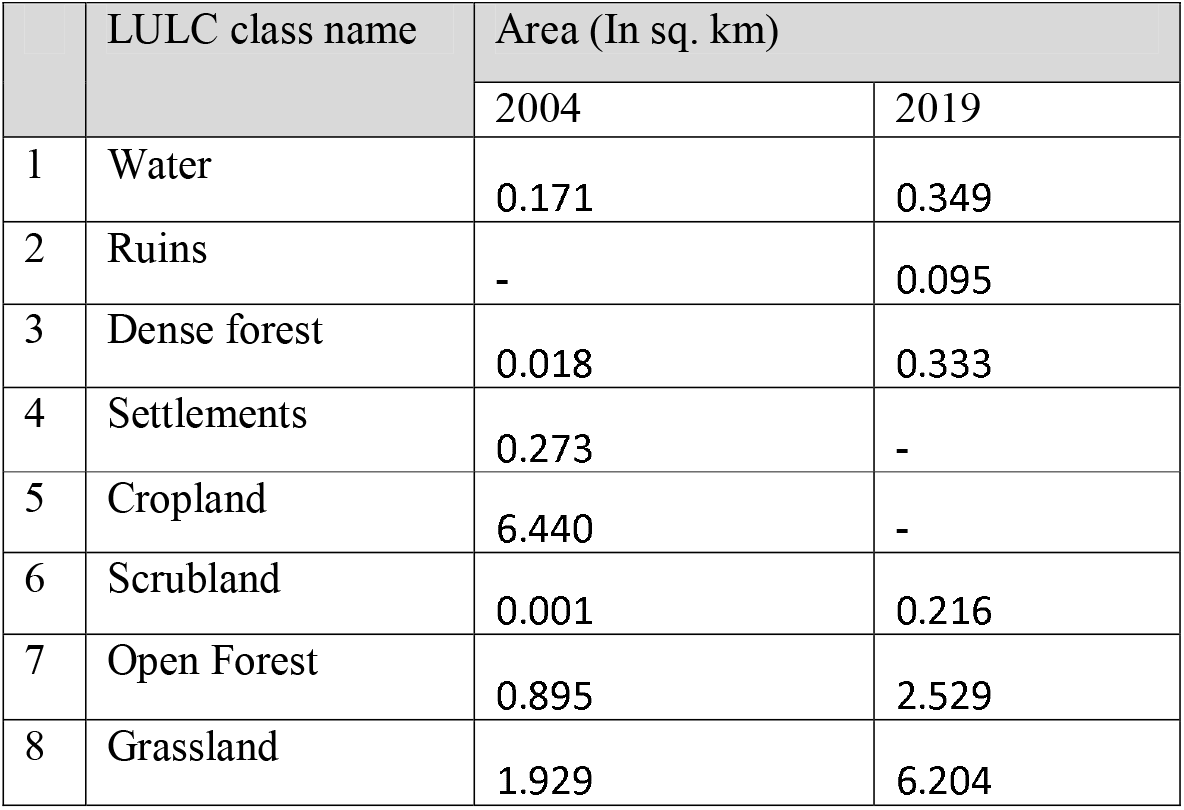
Total area of different LULC classes occupied by studied villages.

|  | LULC class name | Area (In sq. km) |  |
| --- | --- | --- | --- |
|  |  | 2004 | 2019 |
| 1 | Water | 0.171 | 0.349 |
| 2 | Ruins | - | 0.095 |
| 3 | Dense forest | 0.018 | 0.333 |
| 4 | Settlements | 0.273 | - |
| 5 | Cropland | 6.440 | - |
| 6 | Scrubland | 0.001 | 0.216 |
| 7 | Open Forest | 0.895 | 2.529 |
| 8 | Grassland | 1.929 | 6.204 |

### Change matrices

Change matrices were generated between the LULC maps of 2004 and 2019. We found substantial changes (Table 3,4) in the LULC classes from 2004 to 2019. 91.67% of the ruins (empty settlements) are already converted into grasslands. Rest 8.33% of existing ruins are those villages where relocation happened lately in 2009. 10.92% of the grassland has been converted to open forests. The villages’ croplands have converted majorly into grasslands (64.35%) and open forests (29.49%).

**Table 3:** Change matrix of percentage (%) of total area between LULC classes of 2004 (Z) and 2019 (C).

|  | Water<br>(Z1) | Ruins<br>(Z2) | Dense<br>forest<br>(Z3) | Settlements<br>(Z4) | Cropland<br>(Z5) | Scrubland<br>(Z6) | Open<br>forest<br>(Z7) | Grassland<br>(Z8) |
| --- | --- | --- | --- | --- | --- | --- | --- | --- |
| Water (C1) | 63.64% | 0.00% | 0.00% | 0.00% | 1.40% | 0.00% | 1.80% | 6.18% |
| Ruins (C2) | 0.00% | 8.33% | 0.00% | 20.53% | 0.00% | 0.00% | 0.00% | 0.00% |
| Grassland<br>(C3) | 19.25% | 91.67% | 0.00% | 58.61% | 64.35% | 0.00% | 41.92% | 77.77% |
| Open forest<br>(C4) | 9.09% | 0.00% | 63.16% | 11.26% | 29.49% | 0.00% | 42.34% | 10.92% |
| Dense forest<br>(C5) | 7.49% | 0.00% | 36.84% | 6.95% | 2.75% | 0.00% | 9.71% | 1.77% |
| Scrubland<br>(C6) | 0.53% | 0.00% | 0.00% | 2.65% | 1.74% | 0.00% | 4.22% | 2.68% |

**Table 4:** Change matrix of area (in hectare) between LULC classes of the year 2004 and 2019.

|  | **Water**<br>**(Z1)** | **Ruins**<br>**(Z2)** | **Dense**<br>**forest**<br>**(Z3)** | **Settlements**<br>**(Z4)** | **Cropland**<br>**(Z5)** | **Scrubland**<br>**(Z6)** | **Open**<br>**forest**<br>**(Z7)** | **Grassland**<br>**(Z8)** |
| --- | --- | --- | --- | --- | --- | --- | --- | --- |
| **Water (C1)** | 10.705 | 0.00 | 0.00 | 0.00 | 8.906 | 0.00 | 1.529 | 11.604 |
| **Ruins (C2)** | 0.00 | 0.180 | 0.00 | 5.577 | 0.00 | 0.00 | 0.00 | 1.259 |
| **Grassland**<br>**(C3)** | 3.238 | 1.979 | 0.00 | 15.922 | 409.841 | 0.00 | 35.713 | 145.999 |
| **Open forest**<br>**(C4)** | 1.529 | 0.00 | 1.079 | 3.059 | 187.829 | 0.00 | 36.072 | 20.510 |
| **Dense forest**<br>**(C5)** | 1.259 | 0.00 | 0.630 | 1.889 | 17.541 | 0.00 | 8.276 | 3.328 |
| **Scrubland**<br>**(C6)** | 0.090 | 0.00 | 0.00 | 0.720 | 11.065 | 0.00 | 3.598 | 5.038 |

## 4. Discussion

Succession in these villages in different seral stages is prominent. There was a significant increase in Grassland, Open Forest, and Dense Forest from 2004 to 2019, and in areas where the initial relocation happened, ruins are already converted into different forest classes. Higher natural vegetation coverage was expected in the villages relocated initially (Badgadi, Khamariya, Kharaiya) (Figure 2), compared to the others, as these villages were exposed to a natural restoration process for a longer duration. However, no significant increase in the forest classes was found in these three villages. Substantial increase in forest classes in five village (Figure 3) sites: Surajpura, Pipartola, Kaneri, Raipura and Motachaukhan. Although the relocation of these villages occurred between 2007 to 2009, they have more forest types compared to the villages like Badgadi, Khamariya, and Kharaiya, which got relocated from 1987 to 1991. Further research on soil type and nutrition and other environmental factors governing vegetation succession and detailed secondary data on these villages’ early ecological and vegetation structure are needed to draw any conclusion regarding such differences.

A total of 159 plant species, 22 mammalian species (Including Tiger, Jackal, Hyena, Small Indian civet, Sloth bear etc.), and 90 avifauna species were recorded from the relocated village sites (WII, 2022). The recorded species diversity in these vacated villages indicates the success of village relocation in PTR. Feral cattle abandoned by the villagers was documented in 8 of these vacated villages (WII, 2022).

The villages (both those that have been vacated and those that are proposed to be relocated) have been subjected to intense grazing and anthropogenic interferences, such as fire. As a result, once vacated, these villages show the effects of these issues and are frequently infested with invasive weeds. It is not enough to vacate the areas without implementing management interventions over time to ensure resilience capacity, and it is unknown when the resilience capacity will be restored because there are site-specific responses. However, these vacated habitats can become excellent grassland, offering a mosaic of grassland and woodland in the larger context.

In vacated sites, *Cassia sp, Parthenium hirsutus, Hyptis solvencies, Datura sp*., and *Argemone mexicana* are some of the yearly weeds which need attention every year. Lantana is a dominant weed spreading in these areas which needs careful handling as it offers good cover for tigers. Tree growth, mainly, Teak (*Tectona grandis*) and Palash (*Butea monosperma*), is vacated village sites and needs to be monitored for proper management to maintain the vacated village sites at the seral stage of grass meadows.

Under the umbrella of Joint Forest Management/ Ecodevelopment or Social Forestry / Agroforestry/ MGNREGA schemes, grassland development can be carried out in selected villages where land is available. Such areas can be developed into grasslands for wild species and pasture lands for livestock grazing. Grasslands will be restricted for livestock grazing through fencing. The area of pastureland to be developed can only be determined by knowing the cattle population in the surrounding villages. It must be ensured that pastureland supports the carrying capacity of livestock.

## 5. Conclusion

Despite various environmental and forest protection laws, PAs are still vulnerable to anthropogenic pressures. Even if there are no settlements inside the forest, secondary influences of the human population and proper management plans are always required to balance these factors for effective conservation practices. RS-GIS via LULC classification is an essential tool to assess any landscape for repeated intervals and create a baseline for on-ground surveys. We identified the current status of the 14 translocated villages from the Panna tiger reserve and found that succession in different sites varied differently. Besides the change detection, studies like this are essential to identify focus areas within a PA for defining future management strategies and supplementing field surveys. Also, grassland has a significant role in any Central Indian Forest as they support a wide range of taxonomic diversity. Our analysis showed that all these sites now support important areas covering grassland. This research can be helpful for proper monitoring of these grasslands for future management strategies.

